# Learning at variable attentional load requires cooperation between working memory, meta-learning and attention-augmented reinforcement learning

**DOI:** 10.1101/2020.09.27.315432

**Authors:** Thilo Womelsdorf, Marcus R. Watson, Paul Tiesinga

**Author notes:** **Corresponding Authors**: Thilo Womelsdorf and Paul Tiesinga.

## Abstract

Flexible learning of changing reward contingencies can be realized with different strategies. A fast learning strategy involves using working memory of recently rewarded objects to guide choices. A slower learning strategy uses prediction errors to gradually update value expectations to improve choices. How the fast and slow strategies work together in scenarios with real-world stimulus complexity is not well known. Here, we disentangle their relative contributions in rhesus monkeys while they learned the relevance of object features at variable attentional load. We found that learning behavior across six subjects is consistently best predicted with a model combining (*i*) fast working memory (*ii*) slower reinforcement learning from differently weighted positive and negative prediction errors, as well as (*iii*) selective suppression of non-chosen feature values and (*iv*) a meta-learning mechanism adjusting exploration rates based on a memory trace of recent errors. These mechanisms cooperate differently at low and high attentional loads. While working memory was essential for efficient learning at lower attentional loads, enhanced weighting of negative prediction errors and meta-learning were essential for efficient learning at higher attentional loads. Together, these findings pinpoint a canonical set of learning mechanisms and demonstrate how they cooperate when subjects flexibly adjust to environments with variable real-world attentional demands.

**Significance statement:** Learning which visual features are relevant for achieving our goals is challenging in real-world scenarios with multiple distracting features and feature dimensions. It is known that in such scenarios learning benefits significantly from attentional prioritization. Here we show that beyond attention, flexible learning uses a working memory system, a separate learning gain for avoiding negative outcomes, and a meta-learning process that adaptively increases exploration rates whenever errors accumulate. These subcomponent processes of cognitive flexibility depend on distinct learning signals that operate at varying timescales, including the most recent reward outcome (for working memory), memories of recent outcomes (for adjusting exploration), and reward prediction errors (for attention augmented reinforcement learning). These results illustrate the specific mechanisms that cooperate during cognitive flexibility.

## Introduction

Cognitive flexibility is realized through multiple mechanisms (Dajani and Uddin, 2015), including recognizing when the environmental demands change, the rapid updating of expectations and the shifting of response strategies to away from irrelevant towards newly relevant information. The combination of these processes is a computational challenge as they operate on different time scales ranging from slow integration of reward histories to faster updating of expected values given immediate reward experiences (Botvinick et al., 2019). How fast and slow learning processes cooperate to bring about efficient learning is not well understood.

Fast adaptation to changing reward contingencies depends on a fast learning mechanism. Previous studies suggest that such a fast learning strategy can be based on different strategies. One strategy involves memorizing successful experiences in a working memory (WM) and guiding future choices to those objects that have highest expected reward value in working memory (Collins and Frank, 2012; Collins et al., 2014; Viejo et al., 2018; McDougle and Collins, 2020). This WM strategy is similar to recent ‘episodic’ learning models that store instances of episodes as a means to increase learning speed when similar episodes are encountered (Gershman and Daw, 2017; Botvinick et al., 2019).

A second fast learning mechanism uses an attentional strategy that enhances learning from those experiences that were selectively attended (Niv et al., 2015; Rombouts et al., 2015; Oemisch et al., 2019). The advantages of this strategy is an efficient sampling of values when there are many alternatives or uncertain reward feedback (Kruschke, 2011; Farashahi et al., 2017a; Leong et al., 2017). Empirically, such an attentional mechanism accounts for learning values of objects and features within complex multidimensional stimulus spaces (Wilson and Niv, 2011; Niv et al., 2015; Hassani et al., 2017; Leong et al., 2017). In these multidimensional spaces, learning from sampling all possible object instances can be impractical and slows down learning to a greater extent than what is observed in humans and monkeys (Farashahi et al., 2017a; Oemisch et al., 2019). Instead, learners appear to speed up learning by learning stronger from objects that are attended and actively chosen, while penalizing features associated with non-chosen objects (Wilson and Niv, 2011; Niv et al., 2015; Hassani et al., 2017; Leong et al., 2017; Oemisch et al., 2019).

In addition to WM and attention-based strategies, various findings indicate that learning can be critically enhanced by selectively increasing the rate of exploration during difficult or volatile learning stages (Khamassi et al., 2015; Soltani and Izquierdo, 2019). Such a meta-learning strategy, for example, increases the rate of exploring options as opposed to exploiting previously learned value estimates (Tomov et al., 2020). This and other meta-learning approaches have been successfully used to account for learning rewarded object locations in monkeys (Khamassi et al., 2015) and for speeding up learning of multi-arm bandit problems (Wang et al., 2018).

There is evidence for all three proposed strategies in learning, but only a few empirical studies characterize the contribution of different learning strategies. Thus, it is unknown whether working memory, attention-augmented reinforcement learning (RL) and meta-learning approaches are all used and cooperate during learning in differently complex environments.

To address this issue, we set out to test and disentangle the specific contribution of various computational mechanisms for flexibly learning the relevance of visual object features. We trained six monkeys to learn the reward value of object features in environments with varying numbers of irrelevant distracting feature dimensions. Using a larger number of distracting features increased attentional load, resulting in successively slower learning behavior. We found that across monkeys, learning speed was best predicted by a computational RL model that combines working memory, attention augmented RL and meta-learning. The contribution of these individual learning mechanisms varied systematically with attentional load. WM contributed to learning speed particularly at low and medium load, meta-learning contributed at medium and high loads, while selective attention was an essential learning mechanism at all attentional loads.

## Materials and Methods

### Experimental Design

Six male macaque monkeys performed the experiments with an age ranging from 6-9 years and weighting 8.5-14.4 kg. All animal and experimental procedures were in accordance with the National Institutes of Health Guide for the Care and Use of Laboratory Animals, the Society for Neuroscience Guidelines and Policies, and approved by the Vanderbilt University Institutional Animal Care and Use Committee.

The experiment was controlled by USE (Unified Suite for Experiments) using the Unity 3D game engine for behavioral control and visual display (Watson et al., 2019b). Four animals performed the experiment in cage-based touchscreen Kiosk Testing Station described in (Womelsdorf et al., in preparation), while two animals performed the experiment in a sound attenuating experimental booth. All experiments used three dimensionally rendered objects, so called Quaddles (Watson et al., 2019a), that were defined by their body shape, arm style, surface pattern, and color (**Fig. 1A**). We used up to nine possible body shapes, six possible colors, eleven possible arm types and nine possible surface patterns as feature values. The six colors were equidistant within the perceptually defined color space CIELAB. Objects extended ~3cm on the screen corresponding to ~2.5° degrees of visual angle and were presented either on a 24’’ BenQ monitor or an Elo 2094L 19.5 LCD touchscreen running at 60 Hz refresh rate with 1920 × 1080 pixel resolution.

**Figure 1.**
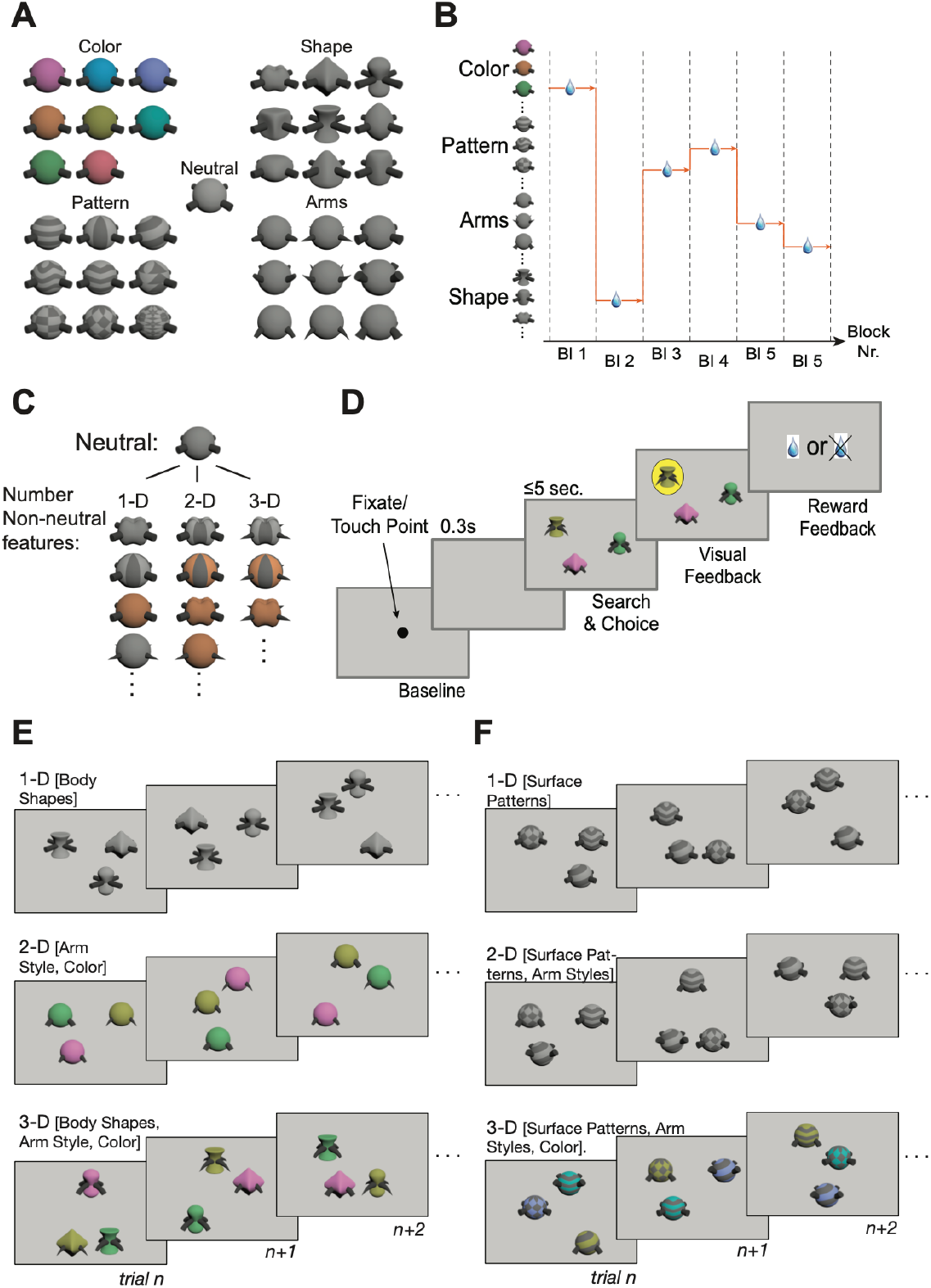
Task paradigm and feature space. (**A**) The task used 3D rendered ‘Quaddle’ objects that varied in color, pattern, shape and arm style. The features grey color, straight arms and spherical body shape were never rewarded in any of the experiments and therefore constitute ‘neutral’ features. (**B**) For successive blocks of 30-60 trials a single feature was rewarded. (**C**) The attentional load conditions differed in the number of non-neutral feature dimensions that varied across trials in a block. Blocks with 1-D, 2-D, and 3-D objects contained stimuli varying features in 1, 2, and 3 features dimensions. (**D**) Trials were initiated by touching or fixating a central stimulus. Three objects were shown at random locations and subjects had to choose one by either touching (four monkeys) or fixating an object (two monkeys) for ≥0.7 sec. Visual feedback indicated correct (yellow) vs error (cyan, not shown) outcomes. Fluid reward followed correct outcomes. (**E**) Sequences of three example trials for a block with 1-D objects (upper row, shape varied), 2-D objects (middle, color and arms varied), and 3-D objects (bottom row, body shape, arms and color) objects. (**F**) Same as *E* but for an object set varying surface pattern, arms, and color.

### Task paradigm

Animals performed a feature-based choice task that required learning through trial-and-error which feature of multidimensional objects is associated with reward. The feature that was reward associated stayed constant for blocks of 35-60 trials and then switched randomly to another feature (**Fig. 1B**). Individual trials (**Fig. 1C**) were initiated by either touching a central blue square (4 monkeys) or fixating the blue square for 0.5 sec. Following a 0.3 sec. delay three objects were presented at random locations of a grid spanning 15 cm on the screen (~24°). The animals had up to 5 sec. to choose one object by touching it for 0.1 sec (four monkeys) or maintaining gaze at an object for 0.7 sec (two monkeys). Following the choice of an object visual feedback was provided as a colored disk behind the selected object (yellow/grey for rewarded/not rewarded choices, respectively) concomitant with auditory feedback (low/high pitched sound for non-rewarded/rewarded choices, respectively). Choices of the object with the rewarded feature resulted in fluid reward at 0.3 sec. following visual and auditory feedback.

For each learning block a unique set of objects was selected that varied in one, two or three feature dimensions from trial-to-trial. The non-varying features were always either a spherical body shape, straight arms with blunt ending, grey color and uniform surface. These feature values were never associated with reward during the experiment and thus represent reward-neutral features. These neutral features defined a neutral object to which we added either one, two, or three non-neutral feature values rendering them 1-, 2-, and 3- dimensional (**Fig. 1D**). For blocks with objects that varied in one feature dimension (1D attentional load condition) three feature values from that dimensions were chosen at the beginning of the block (e.g. body shapes that were oblong, pyramidal, and cubic). One of these features were associated with reward while the two remaining features were not reward-associated, and thus served as distracting features. Within individual trials objects never had the same feature values for these dimensions as illustrated for three successive example trials in **Fig. 1E,F** (*upper* row). The feature values of the unused dimensions were the features of the neutral objects in all trials of that block. For blocks with objects varying in two feature dimensions, a set of three feature values per dimension was selected to obtain nine unique objects combining these features. Only one of the features was associated with reward while the other two feature values of that dimension and the feature values of the other dimension were not linked to reward. **Fig. 1E,F** (*middle* row) illustrates three example trials of these blocks of the 2D attentional load condition. For blocks with objects varying in three feature dimensions (3D attentional load condition), three feature values per dimensions were selected so that the three presented objects had always different features of that dimension which led to twenty-seven unique objects combining these features. Again, only one feature was associated with reward while all other feature values were not linked to reward.

Blocks with objects that varied in 1, 2 and 3 feature dimensions constitute 1D, 2D, and 3D attentional load conditions because they vary the number of feature dimensions that define the search space when learning which feature is rewarded. The specific dimension, feature value, and the dimensionality of the learning problem varied pseudo-randomly from block to block. During individual experimental sessions, monkeys performed up to 30 learning blocks.

### Gaze control

For two animals gaze was monitored with a Tobii Spectrum with 600 Hz sampling rate and binocular infrared eye-tracker. For these animals the experimental session began with a 9-point eye-tracker calibration routine and later reconstruction of object fixations using a robust gaze classification algorithm described elsewhere (Voloh et al., 2020).

### Statistical analysis

All analysis was performed with custom MATLAB code (Mathworks, Natick, MA). Significance tests control for the false discover rate (FDR) with an alpha value of 0.05 to account for multiple comparisons (Benjamini and Hochberg, 1995).

### General formulation of Rescorla-Wagner Reinforcement Learning Models

The value of feature *i* in trial *t*, before the outcome was known, is denoted by 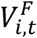. The superscript F stands for feature, in order to distinguish it from the value of an object that will be introduced in the next section. The new value, 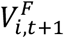 available for decisions on the next trial depends on which features were at t+1 present in the chosen object, and whether this choice was rewarded *R*_*t*_ = 1, or not *R*_*t*_ = 0. The value of features that were present in objects that were not chosen, and those that could appear in the course of the session, but were not present on the current trial, decay with a decay constant 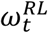, where RL denotes that the decay component is from the reinforcement component of the model as opposed to decay of the working memory introduced below. The features that were present in the chosen and rewarded object increase in value, because the reward prediction error, 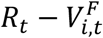 is positive, whereas when the chosen object was not rewarded, the value decays. We have summarized these update rules in the following equations:

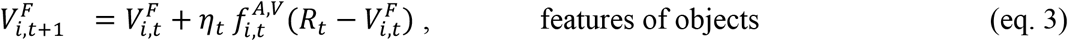

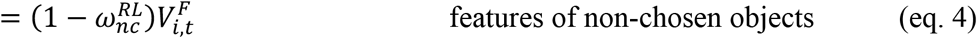

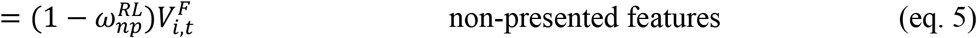

We have indicated a trial-dependence in gain *η* and allow the decay constants ω to depend on whether the feature was present in the non-chosen object (nc) or whether it as part of the stimulus set of the session, but not presented (np) in the current trial. It further carries a superscript RL to indicate it is part of the reinforcement learning formulation rather than working memory (superscript WM). The setting of these parameters depends on the specific model version. In the base RL model, there is no feature-value decay ω_nc,t_ = ω_np,t_ = 0 and the gain is constant and equal to η. In the next model ‘RL gain and loss’, the gain depends on whether the choice was rewarded ηt = *η*_*gain*_ Rt + *η*_*Loss*_ (1 − Rt), which introduces two new parameters *η*_*Gain*_ and *η*_*Loss*_ for rewarded and non-rewarded choices, respectively.

In most models the decay for non-chosen and not presented features was equal ω_nc,t_ = ω_np,t_ = ω^RL^, introducing only a single additional parameter. In the so-called hybrid models we add a feature-dimension gain-factor 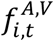, which reflects attention to a particular dimension. It is calculated using a Bayesian model, and is indicated by a superscript V because it affects the value update. Hence, it acts as if information about the role of a certain dimension in the acquisition of the reward is not available. The choice probability 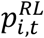 for object *i* at trial *t* is determined using a softmax function:

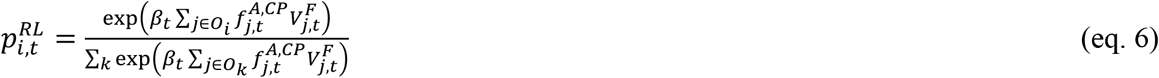

The sum in the exponent of the preceding expression is over the features *j* that are part of object *i*, which defines the set Oi. The factor βt in the exponent determines the extent to which the subject exploits, i.e. systematically chooses the object with the highest compound value, (reflected in a large β) or explores, i.e. makes choices independent of the compound value (reflected in small β). In most model versions the β did not change over trials, while in the meta-learning models with adaptive exploration its value was adaptive, reflecting the history of reward outcomes and is thus trial dependent, see the following subsection.

### Adaptive Exploration

For models with adaptive βt values we follow the model of (Khamassi et al., 2013), which involves determining an error trace:

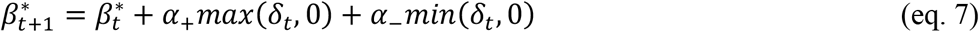

where the min and max functions are used to select the negative and positive part, respectively, of an estimate of the reward prediction error,

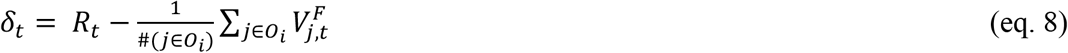

This is a different form of the PE than above, because here we need to consider all features in a chosen object, rather than each feature separately. The error trace is translated into an actual β value using

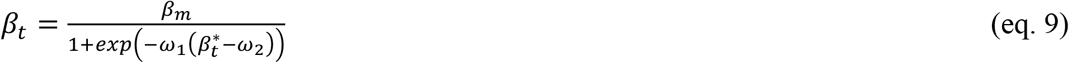

This adaptive sub model replaces one parameter by five new parameters: α+, α−, β_m_, ω_1_ and ω_2_, of which we fixed four in most models, α+ = −0.6, α− = −0.4, ω_1_ = −6, ω_2_ = 0.5, and varied β_m_ and sometimes varied α− as well.

### Attentional dimension weight

The attentional gain factor uses a Bayesian estimate of what the target feature *f* is, hence what the relevant feature dimension is and weighs the contribution of each feature value according to whether it is the target dimension (Niv et al., 2015; Hassani et al., 2017; Oemisch et al., 2019). From the target feature probability 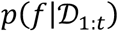 we can obtain a target dimension probability (see (Hassani et al., 2017) for the derivation) by summing over all the feature values *f*(d) that belong to a particular dimension d,

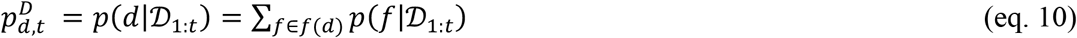

this is turned into a feature gain

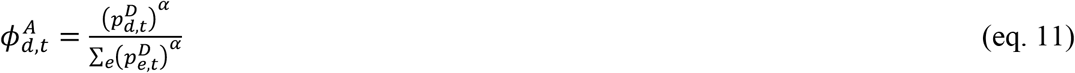

which weighs feature values in each object according to their dimension d(f), for an object *i*, this becomes 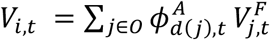, and which we incorporate as feature dependent factor as 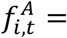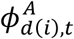 in the relevant expressions (eq. (3) and (6), for the value and choice probability), indicated with additional superscript V and CP, respectively).

### Stickiness in Choice Probability

Stickiness in object choice refers to choosing the object whose feature values overlap with the previously chosen one and represents a kind of perseverance (Balcarras et al., 2016). It is implemented by making the choice probability dependent on whether a feature on the previous trial is present in an object.

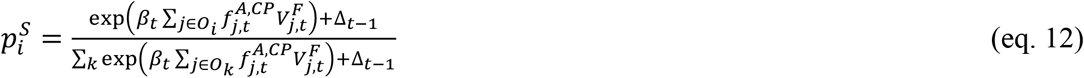

Here Δt−1(i), is equal to e^γ^ −1 when object *i* presented on trial *t* contains at least one feature that was also present in the chosen object on the previous trial (*t* −1). By subtracting one, we assure that when γ = 0, there is no stickiness contribution to the choice. In our setup it is possible that more than one of the current objects contain features that were present in the chosen object.

### Combined Working Memory – Reinforcement models

Working memory models are formulated in terms of the value 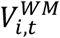 of an object *i* irrespective of what features are present in it (Collins and Frank, 2012). These values are initialized to a non-informative value of 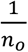, where no is the nouf mobbjeercts, when each of the objects has this value there is no preference in choosing one above the other. When an object is chosen on trial *t*, the value is set to 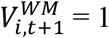 when rewarded, whereas it is reset to the original value 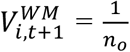 when the choice was not rewarded. All other values decay towards the original value with a decay constant *ω*^*WM*^:

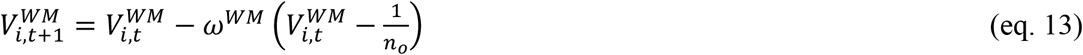

The values are then directly used in the choice probabilities (also denoted pChoice):

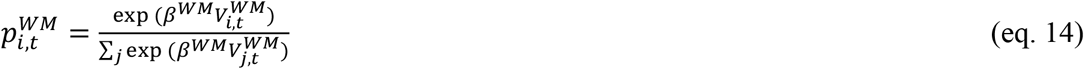

This component mechanism thus introduces two new parameters, a decay constant *ω*^*WM*^ and the softmax parameter *β*^*WM*^, which are separately varied in the fitting procedure.

### Integrating choice probabilities

In the most comprehensive models, choices are determined by a weighted combination of the choice probabilities derived from the RL and WM components, referred to as 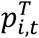 (T stands for total),

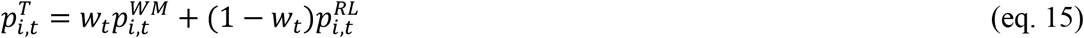

A larger *w*_*t*_ means more weight for the WM predictions in the total choice probability. The update of *w*_*t*_ reflects the value of the choice probability for the choice made and the capacity limitations of the working memory:

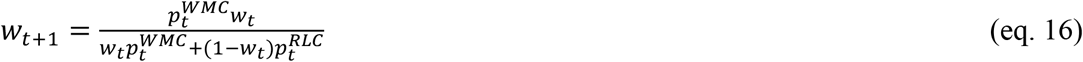

where

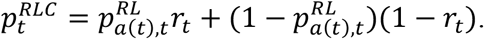

This expression selects from amongst two possible values for 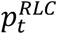 depending on whether *r*_*t*_ = 1 or 0. Here a(*t*) is the index of the object chosen on trial *t*. In addition,

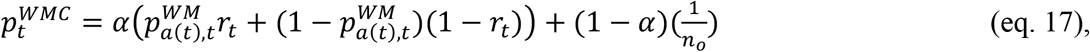

where 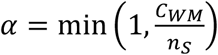 and *C*_*WM*_ is the working memory capacity, essentially the number of objects about which information can be accessed, *n*_*S*_ is the number of objects that can be presented during the task. It is determined as the number of objects whose value 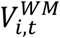 exceeds 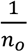 by a margin of 0.01. When n_S_ is much larger than *C*_*WM*_ the information in *p*^*WM*^, which is unlimited in capacity but decays with time, can not be read out, instead 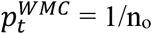. Hence, when 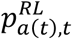 exceeds 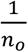 it will win the competition for influence and reduce wt towards WM zero, and with that the influence of 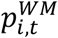.

### Cross-validation

We used a cross-validation procedure for evaluating how well the model predicted the subject’s choices of (test-) learning blocks that were withheld when fitting the model parameters on the remaining (training-) datasets. We repeated the cross-validation 50 times and used the average parameter values across these 50 cross-validation runs to simulate the choices of the monkey. For each cross validation we cut the entire data set at two randomly chosen blocks, yielding three parts. The two largest parts were assigned as training and test set. We did this to keep the trials in the same order as the monkey performed them, as the memory dependent effects in the model (and presumably the monkey) extend beyond the block boundaries. This is different from the standard procedure, where blocks were randomly assigned to test and training sets, hence breaking the block ordering that is important for the model. These model simulations provided the log likelihood (normalized by the number of choices) that reflect how well the model reproduces the monkey’s pattern of correct and erroneous choices. To compare models, we rank-ordered the Akaike Information Criterion (AIC) for each model that penalizes models according to the number of free parameters.

## Results

### Behavioral performance

We measured how six monkeys learned the relevance of object features in a learning task while varying the number of distracting, reward-irrelevant feature dimensions of these objects from one to three. On each trial subjects chose one of three objects and either did or did not receive reward, in order to learn by trial-and-error which object feature predicted reward. The rewarded feature could be any one of 37 possible features values from 4 different feature dimensions (color, shape, pattern and arms) of multidimensional *Quaddle* objects (**Fig. 1A**) (Watson et al., 2019a). The rewarded feature stayed constant within blocks of 35-60 trials (**Fig. 1B**). Learning blocks varied in the number of non-rewarded, distracting features (**Fig. 1C**). Subjects had 5 sec to choose an object which triggered correct or error feedback (a yellow or cyan halo around the chosen object, respectively, **Fig. 1D**). The first of three experimental conditions was labeled 1-dimensional attentional load because all the distractor features were from the same dimension as the target feature (e.g. different body shapes, see examples in top row of **Fig. 1E,F**). At 2-dimensional attentional load, features of a second dimension varied in addition to features from the target feature dimension (e.g. objects varied in body shapes and surface patterning). At 3-dimensional attentional load, object features varied along three dimensions (e.g. varying in body shapes, surface patterns, and arm styles) (bottom row in **Fig. 1E,F**).

Six monkeys performed a total number of 989 learning blocks, completing on average 55/56/54 (SE: 4.4/4.3/4.2; range 41:72) learning blocks for the 1D, 2D, and 3D attentional load conditions, respectively. The number of trials in a block needed to learn the relevant feature, i.e. to reach 75% criterion performance increased for the 1D, 2D, and 3D attentional load condition from on average 6.5, to 13.5, and 20.8 trials (SE’s: 4.2 / 8.3 / 6.9) (Kruskal-Wallis test, p = 0.0152, ranks: 4.8, 10.2, 13.6) (**Fig. 2A,B**). Learning did not differ for blocks with a rewarded feature of the same or of a different dimension as the rewarded feature in the immediately preceding block (intra-versus extradimensional block transitions; Wilcoxon Ranksum test, p = 0.699, ranksum = 36) (**Fig. 2C**). Flexible learning can be influenced by target- and distractor-history effects (Le Pelley et al., 2015; Failing and Theeuwes, 2018; Chelazzi et al., 2019; Rusz et al., 2020), which may vary with attentional load. We first considered the possibility of *latent inhibition* which refers to slower learning of a newly rewarded target feature when that feature was a (learned) distractor in the preceding block than when the target feature was not shown in the previous block. We did, however, not find a latent inhibition effect (paired signed rank test: p = 0.156, signed rank = 3; **Fig. 2D**, *left*). A second history effect is persevering choosing the feature that was a target in the previous block. We quantified this *target perseveration* by comparing learning in blocks in which a previous (learned) target feature became a distractor, to learning blocks in which distractor features were new. We found that target perseveration significantly slowed down learning (paired signed rank test: p = 0.0312, signed rank = 0; **Fig. 2D**, *right*), which was significantly more pronounced in the high (3D) than in the low (1D) attentional load condition (paired signed rank test, again: p = 0.0312, signed rank = 0; **Fig. 2E**). These learning history effects suggest that learned target features had a significant influence on future learning in our task, particularly at high attentional load, while learned distractors had only marginal or no effects on subsequent learning.

**Figure 2.**
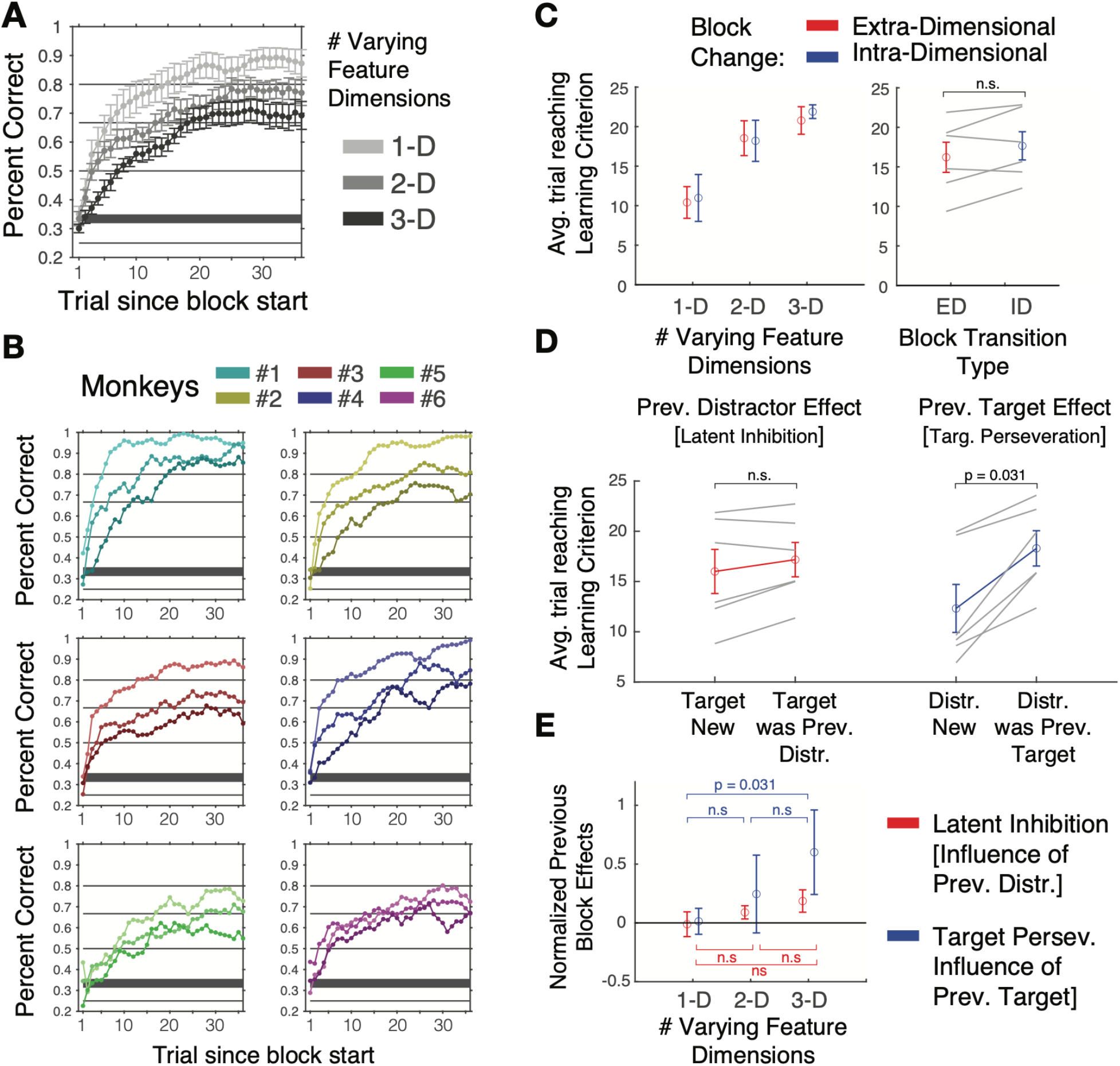
Learning performance. (**A**) Average Learning curves across six monkeys for the 1D, 2D, and 3D load condition. (**B**) Learning curves for each monkeys (colors) for 1D, 2D, 3D (low-to-high color saturation levels). All monkeys showed fastest learning for low load and slowest learning for the high load condition. Curves are smoothed with a 5 trial forward-looking window. (**C**) *Left:* The average trials-to-criterion (75% accuracy over 10 consecutive trials) for low to high attentional load (*x-axis*) for blocks in which the target feature was either of the same - intra-dimensional (ID) - or different dimension - extra-dimensional (ED) - as in the preceding trial. *Right*: Average number of trials-to-criterion across load conditions. Grey lines denote individual monkeys. Errors are SE. (**D**) The red color dneotes average trials-to-criterion for blocks in which the target feature was novel (not shown in previous block), or when it was previously a learned distractor. The blue color denotes the condition in which a distractor feature was either novel (not shown in previous block), or part of the target in the previous block. When distractors were previously targets, learning was slower. (**E**) Latent inhibition of distractors (red) and target perseveration (blue) at low, medium and high load. Errors are SE.

### Multi-component modeling of flexible learning of feature values

To discern specific mechanisms underlying flexible feature-value learning in individual monkeys we fit a series of reinforcement learning (RL) models to their behavioral choices (*see* Material and Methods). These models formalized individual mechanisms and allowed characterizing their role in accounting for behavioral learning at varying attentional load. We started with the classical Rescorla-Wagner reinforcement learner that uses two key mechanisms: (*i*) The updating of value expectations of features *V*^*F*^ every trial *t* by weighting reward prediction errors (PEs) with a learning gain *η*: 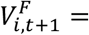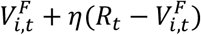 (With reward *R*_*t*_ = 1 for a rewarded choice and zero otherwise), and (*ii*) the probabilistic (‘softmax’) choice of an object *O* given the sum of the expected values of its constituent features *V*_*i*_, 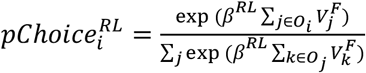 (Sutton and Barto, 2018). These two mechanisms incorporate two learning parameters: the weighting of prediction error (PE) information by *η* (often called the learning rate), *η*(*PE*), and the degree to which subjects explore or exploit learned values represented by *β*, which is small or close to zero when exploring values and larger when exploiting values.

We augmented the Rescorla-Wagner learning model with up to seven additional mechanisms to predict the monkey choices (**Supplementary Table 1**). The first of these mechanisms enhanced the expected values of all object features that were chosen by decaying feature values of non-chosen objects. This selective decay improved the prediction of choices in reversal learning and probabilistic multidimensional feature learning tasks (Wilson and Niv, 2011; Niv et al., 2015; Radulescu et al., 2016; Hassani et al., 2017; Oemisch et al., 2019). It is implemented as decay *ω*^*RL*^ of feature values *V*_*i*_ from non-chosen features and thereby enhanced the value estimate for chosen (and hence attended) features for the next trial *t*:

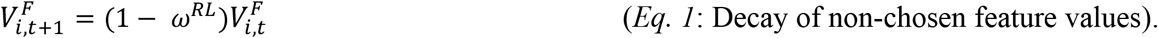

As second mechanism we considered a working memory (WM) process that uploads the identity of rewarded objects in a short-term memory. Such a WM can improve learning of multiple stimulus-response mappings (Collins and Frank, 2012; Collins et al., 2017) and multiple reward locations (Viejo et al., 2015; Viejo et al., 2018). Similar to (Collins and Frank, 2012) we uploaded the value of an object in 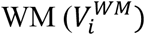 when it was chosen and rewarded and decayed its value with a time constant 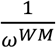. WM proposes a choice using its own choice probability *pChoice*^*WM*^, which competes with the *pChoice*^*RL*^ from the reinforcement learning component of the model. The actual behavioral choice is the weighted sum of the choice probabilities of the WM and the RL component *w*(*pChoice*^*WM*^) + (1 − *w*)*pChoice*^*RL*^. A weight *w* of >0.5 would reflect that the WM content dominates the choice which would be the case when the WM capacity can maintain object values for sufficiently many objects before they fade away (see Methods). This WM module reflects a fast “one–shot” learning mechanism for choosing the recently rewarded object.

As a third mechanism we implemented a meta-learning process that adaptively increases the rate of exploration (the *β* parameter of the standard RL formulation) when errors accumulate. Similar to (Khamassi et al., 2013) the mechanism uses an error trace 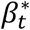, which increases when a choice was not rewarded, by an amount proportional to the negative PE for that choice with a negative gain parameter *α*_−_, and decreases after correct trials proportional to the positive PE weighted by a positive gain parameter *α*_+_ (Khamassi et al., 2013):

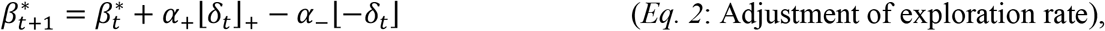

where the PE is given by *δ*_*t*_ − *R*_*t*_ − *V*_*t*_, with V reflecting the mean of all the feature values of the chosen object. The error trace contains a record of the recent reward performance and was transformed into a beta parameter for the softmax choice according to 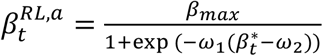 (Khamassi et al., 2013). Transiently increasing the exploration rate increases the chances to find relevant object features when there are no reliable, learned values to guide the choice and there are multiple possible feature dimensions that could be valuable. We kept *α*_+_ = −0.6, *α*_−_ = −0.4 , *ω*_1_ = −6 and *ω*_2_ = 0.5 fixed, and varied *β*_*max*_ and, in some cases, *α*_−_ as well, resulting in a fourth model mechanism that could underlie flexible feature learning under increasing attentional load.

We tested three other neurobiologically plausible candidate mechanisms that played important roles in prior learning studies. A fifth mechanism implemented *choice stickiness* to account for perseverative (repetitive) choices independent of value estimates (Badre et al., 2012; Balcarras et al., 2016). A sixth parameter realized an ‘attentional’ dimension weight during value updates which is realized by multiplying feature values given the reward likelihood for the feature dimension they belong to (Leong et al., 2017; Oemisch et al., 2019). Finally, as a seventh parameter we separately modelled the weighting of negative PE’s after error outcomes, *η*_*Loss*_, and the weighting of positive PE’s for correct outcomes, *η*_*Gain*_, to allow separate learning speeds for avoiding objects that did not lead to reward (after negative feedback) and for facilitating choices to objects that led to rewards (after positive feedback) (Frank et al., 2004; Frank et al., 2007; Kahnt et al., 2009; van den Bos et al., 2012; Caze and van der Meer, 2013; Lefebvre et al., 2017; Taswell et al., 2018). We constructed models that combined two, three or four of these mechanisms. This led to models with two to eight free parameters (*see* Methods). Each model was cross-validated separately for each attentional load condition and for each individual monkey. We calculated the Akaike Information Criteria (AIC) to rank order the models according to how well they predicted actual learning behavior given the number of free parameters.

### Working memory, adaptive exploration and decaying distractor values supplement reinforcement learning

We found that across monkeys and attentional load conditions the RL model that best predicted monkey’s choices during learning had four non-standard components: (*i*) working memory, (*ii*) non-chosen value decay, (*iii*) adaptive exploration rate, and (*iv*) a separate gain for negative PE’s (*η*_*Loss*_) (**Fig. 3**). This model had the lowest Akaike Information Criterion on average across all six monkeys and was ranked first for four individual monkeys (Monkeys 1,3,4, and 5, **Fig 3A,B**; **Supplementary Table 1** shows the complete list of free model parameters for the rank ordered models). It ranked third and fourth for the other two monkeys (**Fig 3A**). Other models ranked in the top 5 contained a stickiness mechanism to account for perseverative choice tendencies (models ranked 2^nd^ and 3^rd^), used an adaptive exploration mechanism with a fixed parameter (that thereby reduced the number of free parameters, model ranked 4th), or lacked the adaptive exploration mechanism (models ranked 3^rd^ and 5^th^). The top-ranked, most-predictive model reproduced well the learning curves obtained from the monkeys (**Fig. 3C**).

**Figure 3.**
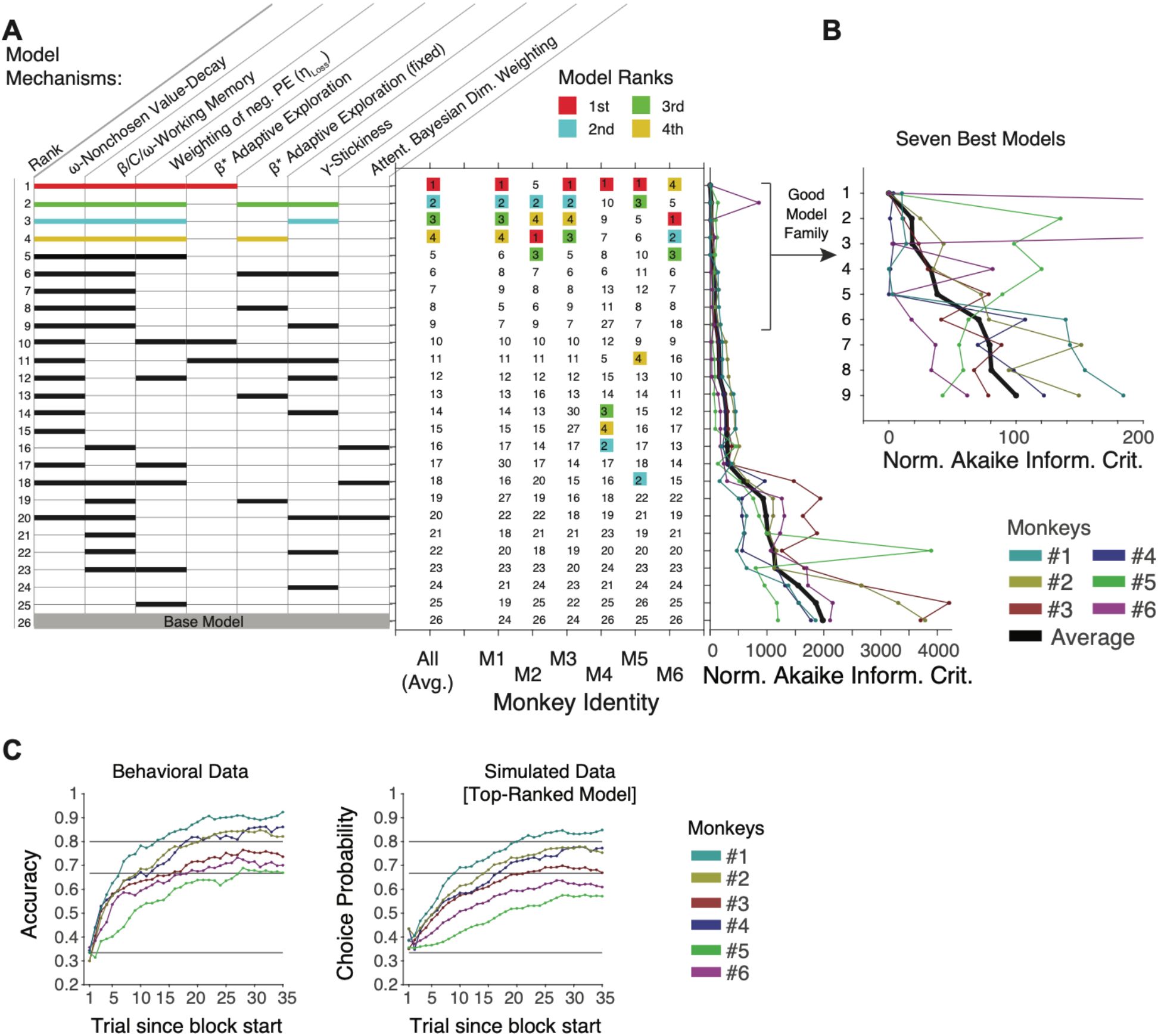
Rank ordering of models with different combinations of mechanisms. (**A**) Models (*rows*) using different combinations of model mechanisms (*columns*) are rank ordered according to their Akaike Information Criterion (AIC). The top-ranked model combined four mechanisms that are highlight in red: Decay of non-chosen features, working memory, adaptive exploration rate, and a separate learning gain for errors (losses). The 2^nd^, 3^rd^, and 4^th^ ranked models are denoted with cyan, green and yellow bars. Thick horizonal bar indicates that the model mechanism was used in that model. The 26^th^ ranked model was the base RL model that used only a beta softmax parameter and a learning rate. *Right*: Model rank average (1^st^ column) and for each individual monkey (columns 2 to 7). See **Supplementary Table 1** for the same table in numerical format with additional information about the number of free parameters for each model. (**B**) After subtracting the AIC of the 1^st^ ranked model, the normalized AIC’s for each monkey confirms that the top-ranked model has low AIC values for each monkey. (**C**) Average behavioral learning curves for the individual monkeys (left) and the simulated choice probabilities of the top-ranked model for each monkey. The simulated learning curves are similar to the monkey learning curves providing face validity for the model.

To discern how the individual model mechanisms of the most predictive model contributed to the learning at low, medium and high attentional load, we simulated the choice probabilities for this full model and for partial models implementing only individual mechanisms of that full model separately for each load condition (**Fig. 4A,B**). This analysis confirmed that the full model was most closely predicting choices of the animals in all load conditions, showing a difference between the model choice probabilities and the monkeys choice accuracy of only ~5% (**Fig. 4C**). The reduced (partial) model that performed similarly well across all attentional loads used the decay of non-chosen features (*ω*^*RL*^) (ranked 16^th^ among all models, **Fig. 4C**). All other partial models were performing differently at low and high attentional loads. The partial working memory model (with *ω*^*WM*^) predicted choices well for the 1D and 2D load conditions but failed to account for choices in the 3D load condition (**Fig. 4C**). The partial model with the adaptive exploration rate (β*) worsened choice probability for the low load condition relative to the standard RL, but improved predictions for the 2D and 3D load condition (**Fig. 4C**). The partial model with the separate weighting of negative prediction errors (*η*_*Loss*_, ranked 25th, see **Fig. 3A**) showed overall better choice probabilities than the standard RL model (ranked 26th), but still failed in predicting 12-14% of the monkeys’ choices when used as the only non-standard RL mechanisms (**Fig. 4C**).

**Figure 4.**
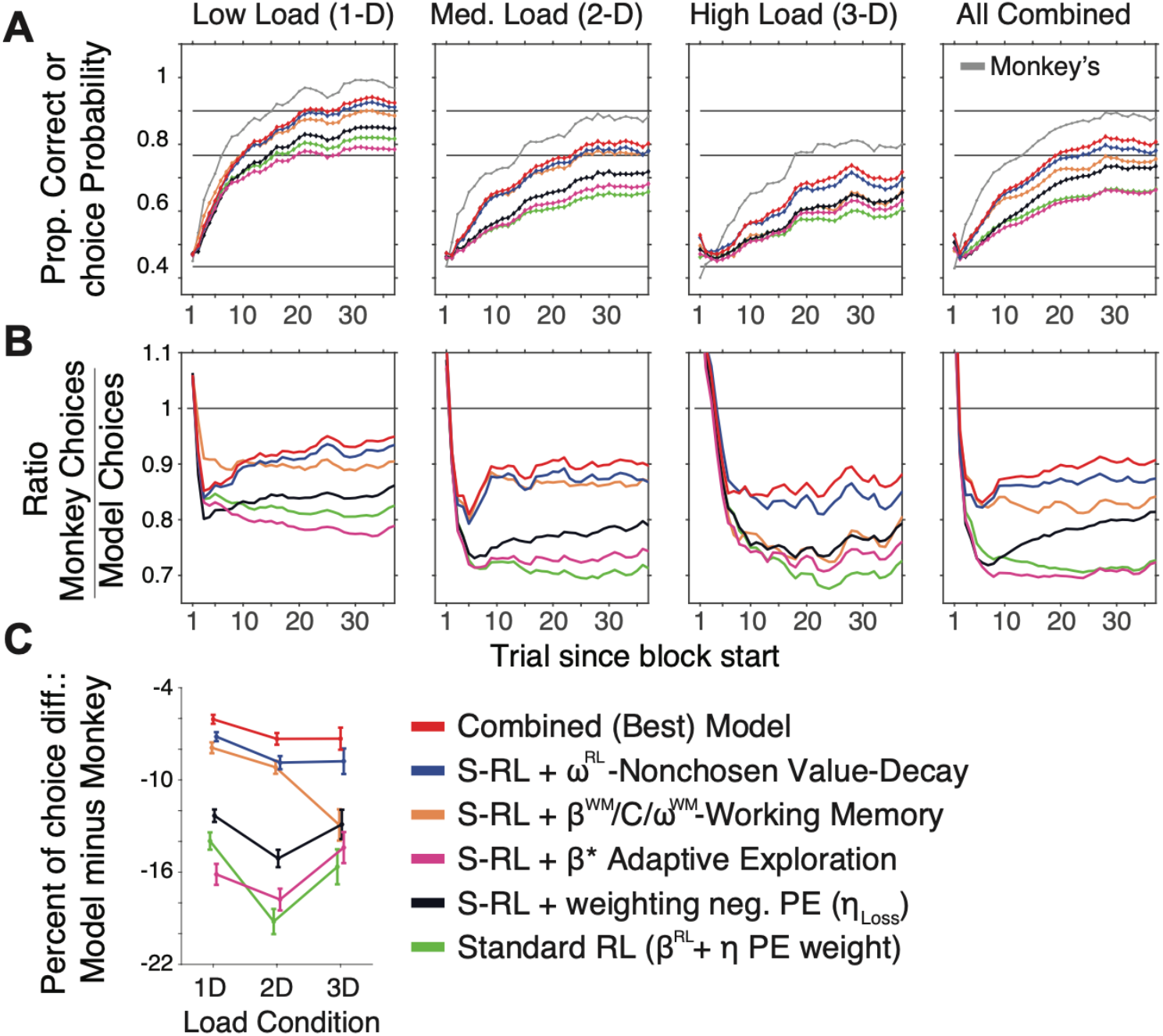
Choice probabilities of monkeys and models at three different loads. (**A**) Average choice accuracy of monkeys (grey) and choice probabilities of six models. The top-ranked model (*red*) combines WM with RL and selective suppression of nonchosen values, a separate learning gain for neg. RPE’s, and adaptive exploration rates. The base RL model (*green*) only contained a softmax beta parameter and a single learning rate. The other models each add a single mechanism to this base model to isolate its contribution to account for the choice patterns of the monkeys. Columns show from left to right the results for low, medium, and high load conditions and for their average. (**B**) The ratio of monkey accuracy and model choice probability shows that in all load conditions, the top-ranked model predicts monkey choices consistently better than models with a single added mechanism. (**C**) Average difference of model predictions (choice probability) and monkeys’ choices (proportion correct) at low to high loads for different models. Error bars are SE.

To understand why WM was only beneficial at low and medium attentional load, but detrimental at high attentional load, we visualized the choice probabilities that the WM module of the full model generated for different objects. We contrasted these WM choice probabilities with the choice probabilities for different stimuli of the RL module and of the combined WM+RL model (**Fig. 5A**). Following a block switch the WM module uploaded an object as soon as it was rewarded and maintained that rewarded object in memory over only few trials. When the rewarded object was encountered again prior to decaying to zero it guided the choice of that object beyond what the RL module would have suggested (evident in trial six in **Fig. 5A-C**). This WM contribution is beneficial when the same object instance re-occurred within few trials, which happened more frequently with low and medium attentional load, but only rarely during high load. At this high load condition, the RL components are faithfully tracking the choice probability of the monkey, while the WM representation of recently rewarded objects is non-informative because (*1*) it can only make a small contribution as the number of stimuli in the block is much larger than the capacity and because (*2*) it does not remember rewarded objects long enough to be around when the objects are presented another time.

**Figure 5.**
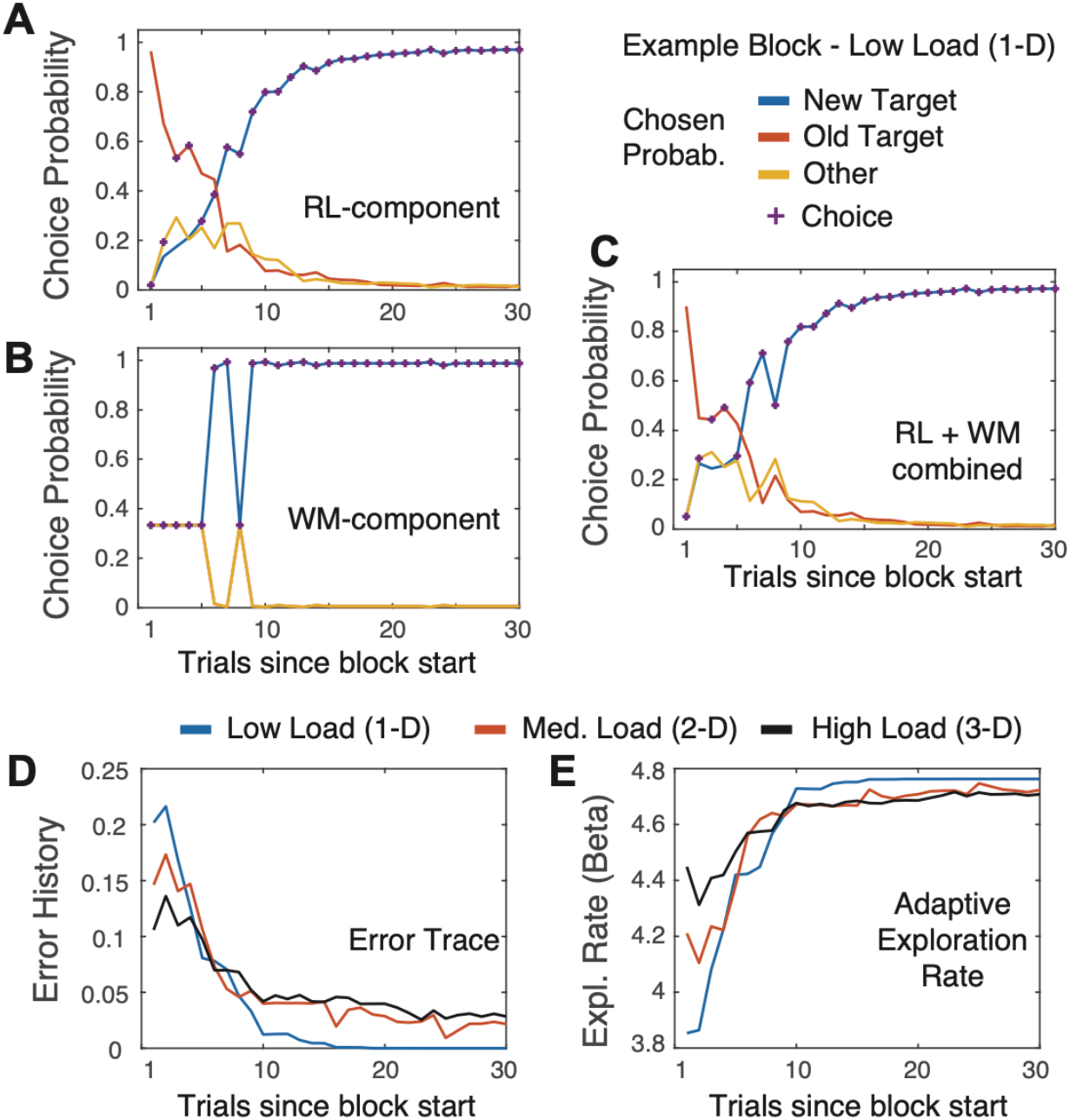
Contribution of working memory, reinforcement learning and adaptive exploration to learning behavior. (**A**) Choice probabilities of the RL component of the top-ranked model for an example block, calculated for the objects with the new target feature (*blue*), the previous block’s target feature (*red*) and other target features (*yellow*). Purple plus signs show which object was chosen. (**B**) Same format and same example block as in *A* but for choice probabilities calculated for objects within the working memory module of the model. Choice probabilities of the WM and the RL component are integrated to reach a final behavioral choice. (**C**) Same as *A* and *B* but after combining the WM and RL components in the full model. Choices closely follow the RL component, but when the WM representation is recently updated its high choice probabilities influences the combined, final choice probability, as evident in trials 6 and 7 in this example block. (**D**) The trace of nonrewarded (error) trials for three example blocks with low, medium and high load peaks immediately after the block switch and then declines for all conditions. Error traces remain non-zero for the medium and high conditions. (**E**) The same example blocks as in *D*. The adaptive exploration rate (y-axis) is higher (lower beta values) when the error trace is high during early trials in a block.

While the WM contribution declined with load, the ability to flexibly adjust exploration rates became more important with high load as is evidenced by improved choice probabilities at high load (**Fig. 4C**). This flexible meta-learning parameter used the trace of recent errors to increase exploration (reflected in lower beta parameter values). Such increases in exploration facilitate disengaging from previously relevant targets after the first errors following the block switch, even when there are no other competitive features in terms of value, because the mechanism enhances exploring objects with previously non-chosen features. Our results suggest that such an adjustment of exploration can reduce the decline in performance at high attentional load (**Fig. 4C**), i.e. when subjects have to balance exploring the increased number of features with acting based on already gained target information (**Fig. 5D,E**).

### The relative contribution of model mechanisms for learning and asymptotic performance

The relative contributions of individual model mechanisms for different attentional loads can be inferred from their load-specific parameter values that best predicted monkey’s learning when cross-validated for learning at each load separately (**Fig. 6**). WM was maintained longer for 2D than 1D (lower *ω*^*WM*^ values), but at 3D load showed fast decay (higher *ω*^*WM*^ values) signifying that WM representations stopped contributing to learning at high load (**Fig. 6C**). When load increased the models showed a consistent decrease of the weighting of positive PE’s (*η*_*Gain*_ from ~0.15 to 0.1) and of the weighting of negative PE’s (*η*_*Loss*_, from ~0.6 to 0.4) (**Fig. 6E**, and **6G**). A potential explanation for the decrease in *η*_*Gain*_ is that with more distracting features more trials are needed to determine what feature is the target, which is achieved with slower updating. The decay of non-chosen feature values (*ω*^*RL*^) was weaker with increased load across monkeys indicating a longer retention of values of non-chosen objects (**Fig. 6F**), which reflects protecting the target value when it is not part of the chosen (hence unrewarded) object. An event that occurs more for high loads. Adaptive exploration rates (*β*\*) increased on average from low to medium and high load (more negative values) signifying increased exploration after errors at higher load.

**Figure 6.**
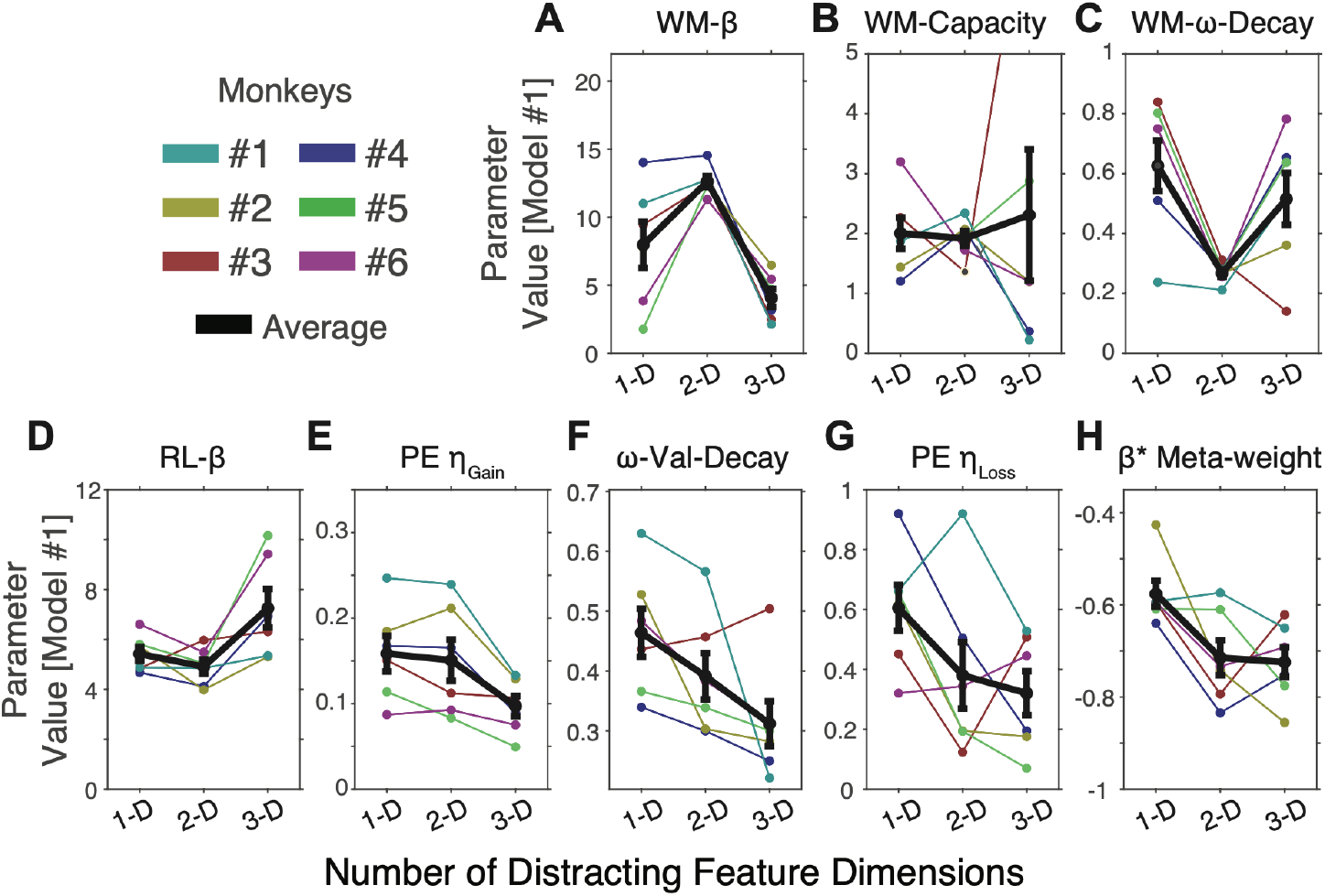
Model parameter values at different attentional loads. The average parameter values (*black*) of the top-ranked model (*y-axis*) plotted against the number of distracting feature dimensions for the WM parameters (**A-C**) and the RL parameters (**D-H**). Individual monkeys are in colors. Error bars indicate SE.

The parameter variations at increasing load could relate either to the learning speed or the asymptotic performance differences at different loads. To quantify their relative contributions, we correlated the values of each model parameter with a model-independent estimate of learning speed (number of trials to reach criterion performance), and with asymptotic accuracy (proportion of correct trials after learning criterion was reached). We found that values of three parameters significantly correlated with learning speed (**Fig. 7A**). Learning occurred significantly earlier (*i*) with larger prediction error weighting for rewarded trials (*η*_*Gain*_, r = −0.69, p = 0.0008, FDR controlled at alpha 0.05), with higher prediction error weight for unrewarded trials (*η*_*Loss*_, r = −0.62, p = 0.0031, FDR controlled at alpha 0.05), and (*iii*) with less decay (and thus better retention) of values of unchosen features (*ω*^*RL*^, r = −0.511, p = 0.0144, 1, FDR controlled at alpha 0.05) (**Fig. 7C**). The same three parameters also correlated significantly with the asymptotic performance that monkey’s showed after learning was achieved (*η*_*Gain*_: r = 0.7, p = 0.0006; *η*_*Loss*_: r = 0.48, p = 0.022; *ω*^*RL*^: r = 0.53, p = 0.0118, 1; all FDR controlled at alpha 0.05). (**Fig. 7B,D**). Higher asymptotic performance was additionally correlated with lower exploration rates (realized by higher β^*RL*^, r = 0.54, p = 0.0097, FDR controlled at alpha 0.05). As for the working memory components, we found that better asymptotic performance was linked to *less* influence of working memory of recently rewarded objects on current choices as reflected in an association of higher *β*^*WM*^ value with better performance (r = 0.54, p = 0.0097, FDR controlled at alpha 0.05). In summary, better performance followed from a combination of choices that exploited value estimates less (lower *β*^*RL*^), kept a stronger value trace for non-chosen features (higher *ω*^*RL*^), and stronger weighting of prediction errors after correct and error outcomes (*η*_*Gain*_, *η*_*Loss*_).

**Figure 7.**
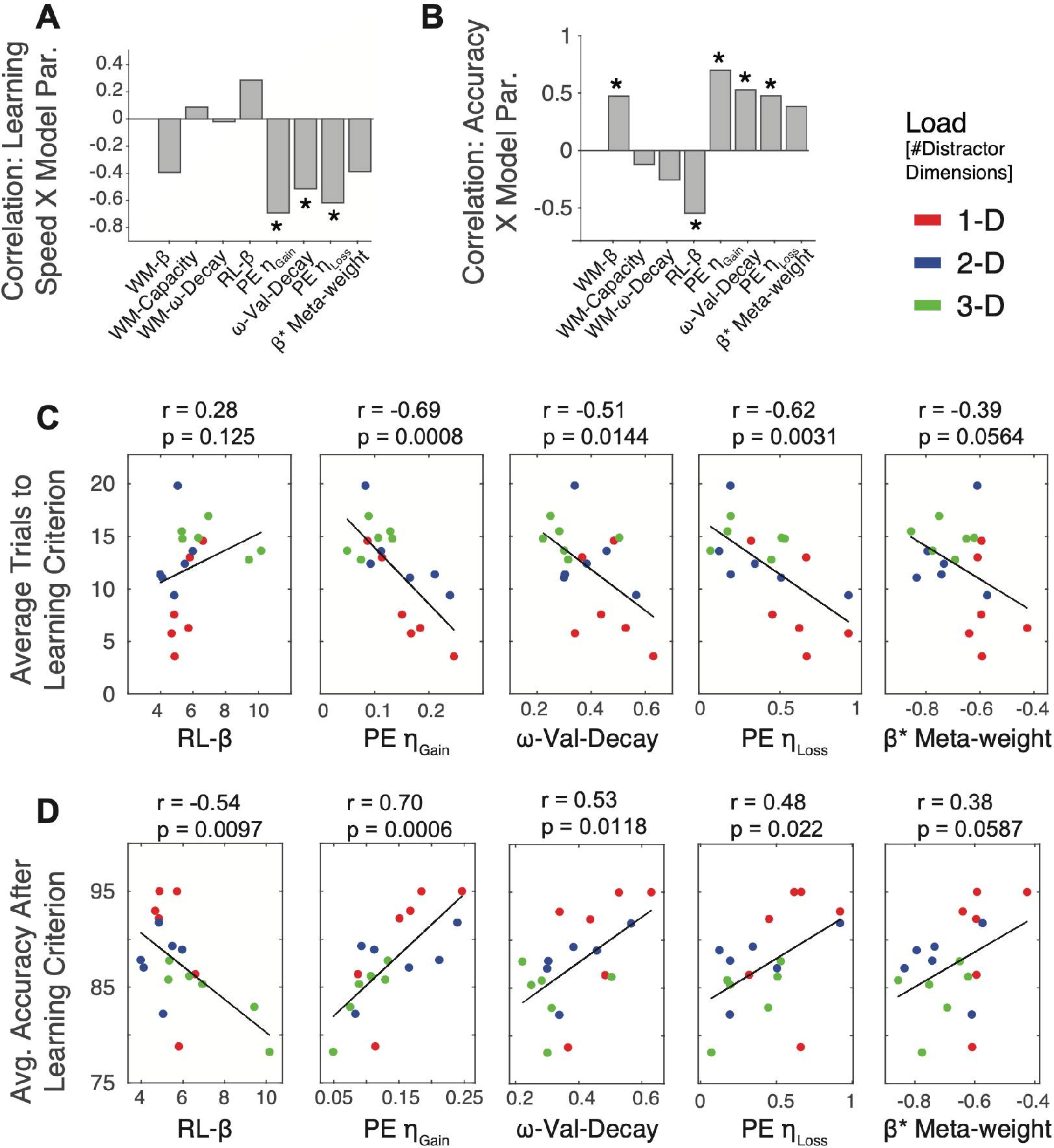
Model parameter values underlying learning speed and asymptotic performance. (A) The correlation across monkeys and attentional load conditions of learning (trials-to-criterion, i.e. less trials signify faster learning) and parameter values of the top-ranked model. Stars denote FDR corrected significance at p < 0.05. Negative correlations denote that higher parameter values associate with faster learning. (**B**) As A but for correlations of monkey’s asymptotic performance accuracy with model parameter values. (**C,D**) Illustration of the correlations shown in *A,B* for the RL parameters. Y-axis shows the learning speed (trial to criterion) and x-axis the parameter values of the top-ranked model. The black line is the regression whose r and p values are given above each plot. Each dot is the average result from one monkey in either the 1D (red), 2-D (blue) or 3-D (green) condition.

### Model parameter values distinguish fast from slow learners

We next tested which model parameters distinguished good from bad learners across attentional load conditions by sorting subjects according to their learning speed (trials-to-criterion). We then correlated the ranks for subjects 1 to 6 for each load condition with the learning speed (**Fig. 8**). Across all parameters the two learning weights (*η*_*Gain*_, *η*_*Loss*_) were significantly larger for subjects that learned faster (*η*_*Gain*_: r = −049, p = 0.019; *η*_*Loss*_: r = −0.51, p = 0.015; significance after control for FDR alpha = 0.05) (**Fig. 8C,D**). This finding illustrates that while multiple mechanism correlate with learning success (**Fig. 7**), the key distinction between good and bad learners lies in their updating speed after errors and after correct outcomes (**Fig. 8**).

**Figure 8.**
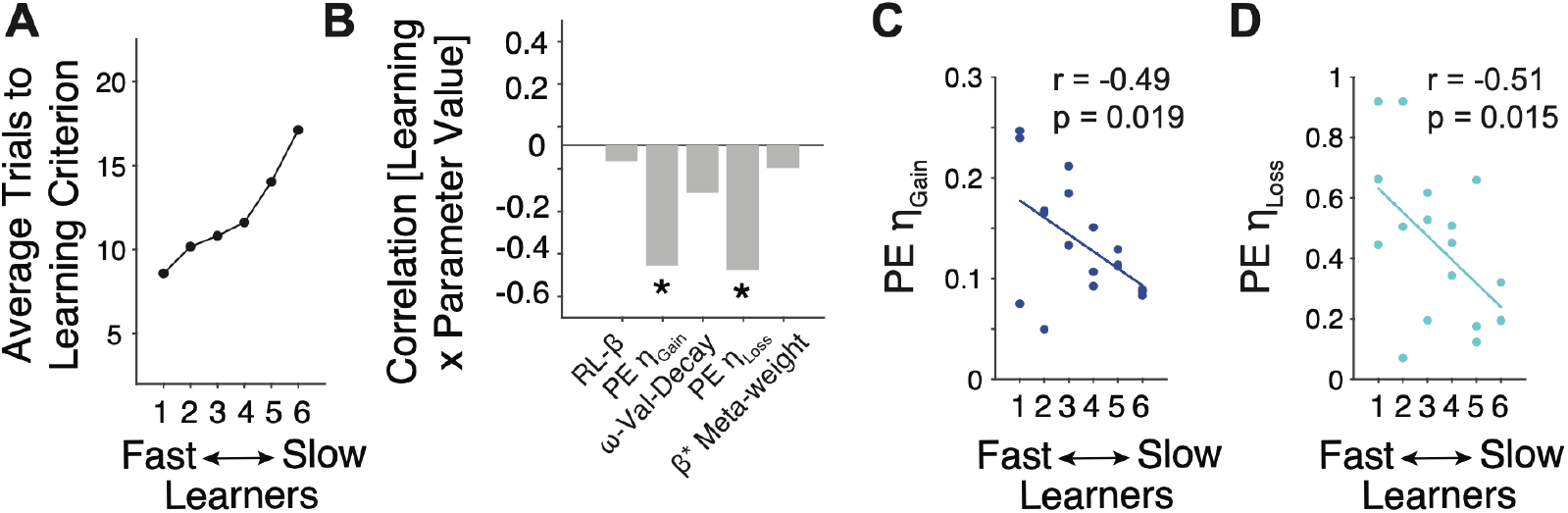
Model mechanisms distinguishing slow and fast learners. (**A**) The average learning speed (the trials to reach criterion, *y-axis*) plotted against the individual monkeys ordered from fastest to slowest learner. (**B**) Correlation of learning speed (trials-to-criterion) and rank-ordered monkeys (ranks #1 to #6). Stars denote FDR corrected significance at p < 0.05. (**C,D**) The weighting of positive prediction errors (*C*) and negative prediction errors (*D*) of the top-ranked model was significantly larger for fast than for slow learners.

## Discussion

We found that learning feature values under increasing attentional load is accounted for by a reinforcement learning framework that incorporates four non-standard RL mechanisms: (*i*) a value-decrementing mechanism that selectively reduces the feature values associated with the non-chosen object, (*ii*) a separate working memory module that retains representations of rewarded objects over a few trials, (*iii*) separate gains for enhancing values after positive prediction errors and for suppressing values after negative prediction errors, and (*iv*) a meta-learning component that adjusts exploration levels according to an ongoing error trace. When these four mechanisms were combined the learning behavior of monkeys was better accounted for than when using fewer or different sets of mechanisms. Critically, the same set of mechanisms were similarly important for all six animals (**Fig. 3**), suggesting they constitute a canonical set of mechanisms underlying cognitive flexibility. Although subjects varied in how these mechanisms were weighted (**Fig. 6**) those with fast learning and hence high cognitive flexibility were distinguished by stronger weighting of positive and negative prediction errors. Taken together these results document a formally defined set of mechanisms that underlies flexible learning of feature relevance under variable attentional load.

### Selective value enhancement is a key mechanism to cope with high attentional load

One key finding was that only one non-standard RL mechanism, the decay of values of non-chosen features (*ω*^*RL*^), contributed similarly to learning across all attentional load conditions, correlating with learning speed and with overall accuracy after learning (r=−0.62, r=0.48, respectively, **Fig 7C,D**). This finding highlights the importance of this mechanism and supports previous studies that used a similar decay of non-chosen features to account for learning in multidimensional environments with deterministic or probabilistic reward schedules (Wilson and Niv, 2011; Niv et al., 2015; Radulescu et al., 2016; Hassani et al., 2017; Oemisch et al., 2019). The working principle of this mechanism is a push-pull effect on the expected values of encountered features and thus resembles a selective attention phenomenon. When a feature is chosen (or attended), its value is updated and contributes to the next choice, while the value of a feature that is not chosen (not attended) is selectively suppressed and contributes less to the next choice. A process with a similar effect has been described in the associability literature whereby the exposure to stimuli without directed attention to it causes a reduction in effective salience of that stimulus. Such reduced effective salience reduces its associability and can cause the latent inhibition of non-chosen stimulus features for learning (Hall and Pearce, 1979; Donegan, 1981; Esber and Haselgrove, 2011) or the slowing of responses to those stimuli (also called negative priming) (Lavie and Fox, 2000). The effect is consistent with a plasticity process that selectively tags synapses of those neuronal connections that represent chosen objects in order to enable their plasticity while preventing (or disabling) plasticity of non-tagged synapses processing non-chosen objects (Rombouts et al., 2015; Roelfsema and Holtmaat, 2018). In computational models such a synaptic tag is activated by feedback connections from motor circuits that carry information about what subjects looked at or manually chose (Rombouts et al., 2015). Accordingly, only chosen objects are updated, resembling how *ω*^*RL*^ implements increasing values for chosen objects when rewarded and the passive decay of values of nonchosen objects. Consistent with this interpretation *ω*^*RL*^ was significantly positively correlated with learning speed (**Fig. 7A**). At low attentional load, high *ω*^*RL*^ values reflected fast forgetting of non-chosen stimuli, while at high attentional load *ω*^*RL*^ adjusted to lower values which slowed down the forgetting of values associated with nonchosen objects. The lowering of the *ω*^*RL*^ decay at high load reflects that values of all stimulus features are retained in a form of choice-history trace. Consistent with this finding, various studies have reported that prefrontal cortex areas contain neurons representing values of unchosen objects and unattended features of objects (Boorman et al., 2009; Westendorff et al., 2016). Our results demonstrate that at high attentional load, the ability of subjects to retain the value history of those nonchosen stimulus features is a critical factor for fast learning and good performance levels (**Fig. 7A**).

### Working memory supports learning in parallel to reinforcement learning

We found empirical evidence that learning the relevance of visual features leverages a fast working memory mechanism in parallel with a slower reinforcement learning of values. This finding empirically documents the existence of parallel (WM and RL) choice systems, each contributing to the monkey’s choice in individual trials to optimize outcomes. The existence of such parallel choice and learning systems for learning fast and slow has a long history in the decision-making literature (Poldrack and Packard, 2003; van der Meer et al., 2012; Balleine, 2019). For example, WM has been considered to be the key component for a rule-based learning system that uses a memory of recent rewards to decide to stay or switch response strategies (Worthy et al., 2012). A separate learning system is associative and implicitly integrates experiences over longer time periods (Poldrack and Packard, 2003), which in our model corresponds to the reinforcement learning module.

The WM mechanisms we adopted for the feature learning task is similar to WM mechanisms that contributed in previous studies to the learning of strategies of a matching pennies game (Seo et al., 2014), the learning of hierarchical task structures (Collins and Frank, 2012, 2013; Alexander and Brown, 2015), or the flexible learning of reward locations (Viejo et al., 2015; Viejo et al., 2018). Our study adds to these prior studies by documenting that the benefit of WM is restricted to tasks with low and medium attentional load. The failure of working memory to contribute to learning at higher load might reflect an inherent limit in working memory capacity. Beyond an interpretation that WM capacity limits are reached at higher load, WM is functionally predominantly used to facilitate processing of actively processed items as opposed to inhibiting the processing of items stored in working memory (Noonan et al., 2018). In other words, a useful working memory is rarely filled with distracting, non-relevant information that a subject avoids. In our task, high distractor load would thus overwhelm the working memory store with information about non-rewarded objects whose active use would not lead to reward. Consequently, the model – and the subject whose choices the model predicts – downregulate the importance of WM at high attentional load, relying instead on slower reinforcement learning mechanism to cope with the task.

### Separate learning rates promote avoiding choosing objects resulting in worse-than-expected outcomes

We found that separating learning from positive and negative prediction errors improved model predictions of learning across attentional loads (**Fig. 4B**) by allowing a ~3-fold larger learning rate for negative than positive outcomes. Thus, monkeys were biased to learn faster to avoid objects with worse-than-expected feature values than to stay with choosing objects with better-than-expected feature values. A related finding is the observation of larger learning rates for losses than gains for monkeys performing a simpler object-reward association tasks (Taswell et al., 2018). In our task, such a stronger weighting of erroneous outcomes seems particularly adaptive because the trial outcomes were deterministic, rather than probabilistic, and thus a lack of reward provided certain information that the chosen features were part of the distracting feature set. Experiencing an omission of reward can therefore immediately inform subjects that feature values of the chosen object should be suppressed as much as possible to avoid choosing it again. This interpretation is consistent with recent computational insights that the main effect of having separate learning rates for positive and negative outcomes is to maximize the contrast between available values for optimized future choices (Caze and van der Meer, 2013). According to this rationale, larger learning rates for negative outcomes in our task promotes switching away from choosing objects with similar features again in the future. We should note that in studies with uncertain (probabilistic) reward associations that cause low reward rates, the overweighting of negative outcomes would be non-adaptive as it would promote frequent switching choices which is suboptimal in these probabilistic environments (Caze and van der Meer, 2013). These considerations can also explain why multiple prior studies with probabilistic reward schedules report of an overweighting of positive over negative prediction errors which in their tasks promoted staying with and prevent switching from recent choices (Frank et al., 2004; Frank et al., 2007; Kahnt et al., 2009; van den Bos et al., 2012; Lefebvre et al., 2017).

The separation of two learning rates also demonstrates that our task involves two distinct learning systems for updating values after experiencing nonrewarded and rewarded choice outcomes. Neurobiologically, this finding is consistent with studies of lesioned macaques reporting that learning from aversive outcomes is more rapid than from positive outcomes and that this rapid learning is realized by fast learning rates in the amygdala as opposed to slower learning rates for better-than-expected outcomes that closely associated with the ventral striatum (Namburi et al., 2015; Averbeck, 2017; Taswell et al., 2018). Our finding of considerably higher (faster) learning rates for negative than positive prediction errors is consistent with this view of a fast versus a slow RL updating system in the amygdala and the ventral striatum, respectively. The importance of these learning systems for cognitive flexibility is evident by acknowledging that learning rates from both, positive and negative outcomes, distinguished good and bad learners (**Fig. 8**), which supports reports that better and worse learning human subjects differ prominently in their strength of prediction error updating signals (Klein et al., 2007; Schonberg et al., 2007; Krugel et al., 2009).

### Adaptive exploration contributes at high attentional load

We found that adaptive increases of exploration during the learning period contributed to improved learning at high load (**Fig. 3**). Adapting the rate of exploration over exploitation reflects a meta-learning strategy that changes the learning process itself by adaptively enhancing searching for new choice options irrespective of already acquired expected value (Doya, 2002). Our finding critically extends insights that adaptive learning rates are critically important to cope with uncertain environments (Farashahi et al., 2017b; Soltani and Izquierdo, 2019) to target uncertainty imposed by increased distractor load. In earlier studies, reward uncertainty was estimated to adjust learning rate in tasks with varying volatility (Farashahi et al., 2017b), changing outcome probabilities when predicting sequences of numbers (Nassar et al., 2010), sharp transitions of exploratory search for reward rules and exploitation of those rules (Khamassi et al., 2015), probabilistic reward schedule during reversal learning (Krugel et al., 2009), or the compensation for error in multi-joint motor learning (Schweighofer and Arbib, 1998). A commonality of these prior meta-learning studies is a relatively high level of uncertainty about the source for reward or error outcomes. In our task, the uncertainty about the target feature systematically increases with the number of distracting features. As a consequence of enhanced uncertainty, subjects utilized a learning mechanism that increased randomly exploring new choice options when non-rewarded choices accumulated, and to reduce exploring alternative choices when choices began to lead to reward outcomes. Such balancing of exploration and exploitation can be achieved by using a memory of recent reward history to adjust undirected vigilance (Dehaene et al., 1998; Khamassi et al., 2013) or other forms of exploratory strategies (Tomov et al., 2020).

## Conclusion

In summary, our study documents that a pure reinforcement learning modeling approach does not capture the cognitive processes needed to solve feature-based learning. By formalizing the subcomponent processes needed to augment standard RL modeling we provide strong empirical evidence for the recently proposed ‘EF-RL’ framework that describes how executive functions (EF) augment RL mechanism during cognitive tasks (Rmus et al., 2020). The framework asserts that RL mechanisms are central for learning a policy to address task challenges, but that attention-, action-, and higher-order expectations are integral for shaping these policies (Rmus et al., 2020). In our study these ‘EF’ functions included working memory, adaptive exploration, and an attentional mechanism for forgetting nonchosen values. As outlined in **Fig. 9** these three mechanisms leverage distinct learning signals, updating values based directly on outcomes (WM), on prediction errors (RL based decay of nonchosen values), or on a continuous error history trace (meta-learning based adaptive exploration). As a consequence, these three learning mechanisms operate in parallel and influence choices to variable degrees across different load conditions, for instance, learning fast versus slow (WM versus Meta-learning versus RL) and adapting optimally to low versus high attentional load (WM versus Meta-learning). Our study documents that these mechanisms operate in parallel when monkeys learn the relevance of features, providing an starting point to identify how neural systems integrate these mechanisms during cognitive flexible behavior.

**Figure 9.**
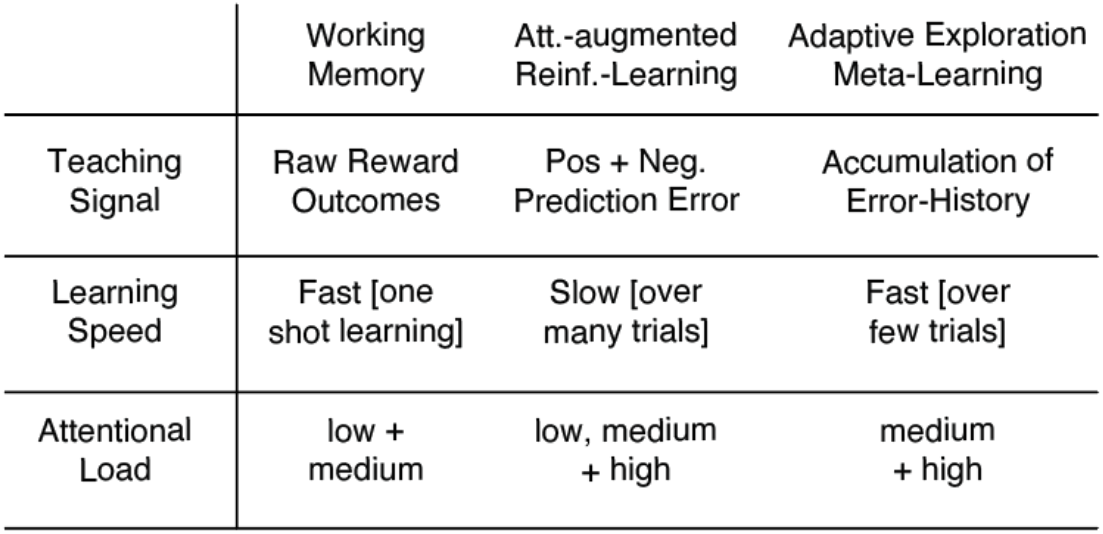
Characteristics of the working memory, reinforcement learning and meta-learning components. The model components differ in the teaching signals that trigger adjustment, in the learning speed and in how important they are to contribute to learning at increasing attentional load.

## Acknowledgements

The authors would like to thank Shelby Volden and Seth König for help with animal training and data collection, and Seth König for preparing the data for analysis and for introducing the concept of a ‘neutral’ Quaddle stimulus. This work was supported by a grant from the National Institute of Biomedical Imaging and Bioengineering of the National Institutes of Health (R01EB028161). The funder had no role in study design, data collection and analysis, the decision to publish, or the preparation of this manuscript.

## Data and code accessibility

Data and computational modeling code for reproducing the results of the best fitting model (**Fig. 4**) is available on a github link that is activated upon publication of this manuscript.

## Author Contributions

T.W. and M.R.W. conceived the experiments. M.R.W. wrote the code to run the experiments. T.W. and P.T. analyzed the data. T.W., M.R.W., and P. T. wrote the paper.

---

**Supplementary Table 1.** Overview of the parameter used in models evaluated and ranked according to AIC higher than the base RL model (which is the model ranked 26th). See figure 3 of the main text for a graphical illustration of the model rank ordering.

| Model Rank | RL gain ($\eta$) | RL decay ($\omega$) | RL softmax ($\beta$) | RL atn ($\alpha$) | WM decay ($\omega$) | WM softmax ($\beta$) | WM capacity ($C_{WM}$) | stickiness ($\gamma$) | #para |
| --- | --- | --- | --- | --- | --- | --- | --- | --- | --- |
| 1 | $\eta_{Gain}, \eta_{Loss}$ | $\omega^{RL}$ | $\beta_m, \alpha_- (\alpha_+, \omega_1, \omega_2 \text{ fix})$ | - | $\omega^{WM}$ | $\beta^{WM}$ | $C_{WM}$ | - | 8 |
| 2 | $\eta_{Gain}, \eta_{Loss}$ | $\omega^{RL}$ | $\beta_m (\alpha_+, \alpha_-, \omega_1, \omega_2 \text{ fix})$ | - | $\omega^{WM}$ | $\beta^{WM}$ | $C_{WM}$ | $\gamma$ | 8 |
| 3 | $\eta_{Gain}, \eta_{Loss}$ | $\omega^{RL}$ | $\beta^{RL}$ | - | $\omega^{WM}$ | $\beta^{WM}$ | $C_{WM}$ | $\gamma$ | 8 |
| 4 | $\eta_{Gain}, \eta_{Loss}$ | $\omega^{RL}$ | $\beta_m (\alpha_+, \alpha_-, \omega_1, \omega_2 \text{ fix})$ | - | $\omega^{WM}$ | $\beta^{WM}$ | $C_{WM}$ | - | 7 |
| 5 | $\eta_{Gain}, \eta_{Loss}$ | $\omega^{RL}$ | $\beta^{RL}$ | - | $\omega^{WM}$ | $\beta^{WM}$ | $C_{WM}$ | - | 7 |
| 6 | $\eta$ | $\omega^{RL}$ | $\beta_m (\alpha_+, \alpha_-, \omega_1, \omega_2 \text{ fix})$ | - | $\omega^{WM}$ | $\beta^{WM}$ | $C_{WM}$ | - | 7 |
| 7 | $\eta$ | $\omega^{RL}$ | $\beta^{RL}$ | - | $\omega^{WM}$ | $\beta^{WM}$ | $C_{WM}$ | - | 6 |
| 8 | $\eta$ | $\omega^{RL}$ | $\beta_m (\alpha_+, \alpha_-, \omega_1, \omega_2 \text{ fix})$ | - | $\omega^{WM}$ | $\beta^{WM}$ | $C_{WM}$ | - | 6 |
| 9 | $\eta$ | $\omega^{RL}$ | $\beta_m, \alpha_- (\alpha_+, \omega_1, \omega_2 \text{ fix})$ | - | $\omega^{WM}$ | $\beta^{WM}$ | $C_{WM}$ | $\gamma$ | 7 |
| 10 | $\eta_{Gain}, \eta_{Loss}$ | $\omega^{RL}$ | $\beta_m, \alpha_- (\alpha_+, \omega_1, \omega_2 \text{ fix})$ | - | - | - | - | - | 5 |
| 11 | $\eta_{Gain}, \eta_{Loss}$ | $\omega^{RL}$ | $\beta_m (\alpha_+, \alpha_-, \omega_1, \omega_2 \text{ fix})$ | - | - | - | - | $\gamma$ | 5 |
| 12 | $\eta_{Gain}, \eta_{Loss}$ | $\omega^{RL}$ | $\beta^{RL}$ | - | - | - | - | $\gamma$ | 5 |
| 13 | $\eta$ | $\omega^{RL}$ | $\beta_m (\alpha_+, \alpha_-, \omega_1, \omega_2 \text{ fix})$ | - | - | - | - | - | 3 |
| 14 | $\eta$ | $\omega^{RL}$ | $\beta^{RL}$ | - | - | - | - | $\gamma$ | 4 |
| 15 | $\eta$ | $\omega^{RL}$ | $\beta^{RL}$ | - | - | - | - | - | 3 |
| 16 | $\eta$ | $\omega^{RL}$ | $\beta^{RL}$ | $\alpha$ | $\omega^{WM}$ | $\beta^{WM}$ | $C_{WM}$ | - | 7 |
| 17 | $\eta_{Gain}, \eta_{Loss}$ | $\omega^{RL}$ | $\beta^{RL}$ | - | | | | - | 4 |
| 18 | $\eta_{Gain}, \eta_{Loss}$ | $\omega^{RL}$ | $\beta^{RL}$ | $\alpha$ | $\omega^{WM}$ | $\beta^{WM}$ | $C_{WM}$ | - | 8 |
| 19 | $\eta$ | - | $\beta_m (\alpha_+, \alpha_-, \omega_1, \omega_2 \text{ fix})$ | - | $\omega^{WM}$ | $\beta^{WM}$ | $C_{WM}$ | - | 5 |
| 20 | $\eta$ | $\omega^{RL}$ | $\beta^{RL}$ | $\alpha$ | $\omega^{WM}$ | $\beta^{WM}$ | $C_{WM}$ | $\gamma$ | 8 |
| 21 | $\eta$ | - | $\beta^{RL}$ | - | $\omega^{WM}$ | $\beta^{WM}$ | $C_{WM}$ | - | 5 |
| 22 | $\eta$ | - | $\beta^{\text{RL}}$ | - | $\omega^{\text{WM}}$ | $\beta^{\text{WM}}$ | $C_{\text{WM}}$ | $\gamma$ | 6 |
| 23 | $\eta_{\text{Gain}}, \eta_{\text{Loss}}$ | - | $\beta^{\text{RL}}$ | - | $\omega^{\text{WM}}$ | $\beta^{\text{WM}}$ | $C_{\text{WM}}$ | - | 6 |
| 24 | $\eta$ | - | $\beta^{\text{RL}}$ | - | - | - | - | $\gamma$ | 3 |
| 25 | $\eta_{\text{Gain}}, \eta_{\text{Loss}}$ | - | $\beta^{\text{RL}}$ | - | - | - | - | - | 3 |
| 26 | $\eta$ | - | $\beta^{\text{RL}}$ | - | - | - | - | - | 2 |

